# The first chromosome-level gecko genome reveals dynamic sex chromosomes in Neotropical leaf-litter geckos (Sphaerodactylidae: *Sphaerodactylus*)

**DOI:** 10.1101/2021.08.13.456260

**Authors:** Brendan J. Pinto, Shannon E. Keating, Stuart V. Nielsen, Daniel P. Scantlebury, Juan D. Daza, Tony Gamble

## Abstract

Sex chromosomes have evolved many times across eukaryotes, indicating both their importance and their evolutionary flexibility. Some vertebrate groups, such as mammals and birds, have maintained a single, conserved sex chromosome system across long evolutionary time periods. By contrast, many reptiles, amphibians, and fish have undergone frequent sex chromosome transitions, most of which remain to be catalogued. Among reptiles, gecko lizards (infraorder Gekkota) have shown an exceptional lability with regard to sex chromosome transitions and may possess the majority of transitions within squamates (lizards and snakes). However—across geckos—information about sex chromosome linkage is expressly lacking, leaving large gaps in our understanding of the evolutionary processes at play in this system. To address this gap, we assembled the first chromosome-level genome for a gecko and use this linkage information to survey six *Sphaerodactylus* species using a variety of genomic data, including whole-genome re-sequencing, RADseq, and RNAseq. Previous work has identified XY systems in two species of *Sphaerodactylus* geckos. We expand upon that work to identify between two and four sex chromosome *cis*-transitions (XY to XY) within the genus. Interestingly, we confirmed two linkage groups as XY sex chromosome systems that were previously unknown to act as sex chromosomes in tetrapods (syntenic with *Gallus* 3 and *Gallus* 18/30/33). We highlight the increasing evidence that most (if not all) linkage groups will likely be identified as a sex chromosome in future studies given thorough enough sampling.

## Introduction

Sexual reproduction is ubiquitous in vertebrates but the ways in which species determine sex can differ (Graves, 2008; Otto and Lenormand, 2002). Most vertebrate species determine sex using genetic cues inherited from one of their parents (i.e. sex chromosomes), either from the sperm (male heterogamety; XY) or the egg (female heterogamety; ZW). Traditionally, cytogeneticists identified sex chromosomes by karyotyping a male and female of a species and looking for morphological differences between the two karyotypes (Stevens, 1905). Until recently, the majority of sex chromosome research was restricted to species whose sex chromosomes were heteromorphic, or visibly different under a light microscope, such as mammals (XY) and birds (ZW). As a consequence, much of what we know about vertebrate sex chromosomes comes from studies in mammals and birds who possess ancient, degenerated sex determination systems (SDS), where transitions are rare or non-existent (Bachtrog, 2003; Graves, 2008; Ohno, 1967; Zhou et al. 2014). However, other vertebrate groups, such as fish, amphibians, and squamate reptiles, typically possess homomorphic sex chromosomes, which appear identical under the light microscope, effectively stifling any investigations of sex chromosome evolution in these groups (Ezaz et al. 2009; Hillis and Green 1990; and reviewed in Gamble, 2010).

Sex chromosomes evolve when one chromosome of an autosomal pair acquires a sex determining allele (Graves, 2008; Ohno 1967). Through shifts in selective pressure between the nascent sex chromosome pair (e.g. sexually antagonistic selection) and chromosomal rearrangements (e.g. inversions), recombination can be suppressed between the X/Y or Z/W chromosomes (Charlesworth, 1991; Ohno, 1967). After recombination is suppressed—with no mechanism to repair deleterious mutations—the sex-limited chromosome (Y or W) begins to accumulate mutations and degenerate by losing functional copies of genes and gaining segments of repetitive DNA (Bull, 1983; Bachtrog, 2013; Charlesworth, 1991; Charlesworth and Charlesworth, 2000; Muller, 1918; 1964; Ohno, 1967; Wright et al. 2016). This non-recombining region slowly expands outward over time generating what are known as “evolutionary strata” (Bachtrog, 2013; García-Moreno and Mindell, 2000; Graves, 2008; Lahn and Page, 1999). In taxa where transitions in sex chromosome systems are frequent, the degree in which species conform to the progressive model of sex chromosome evolution described above can differ. In effect, groups can possess homomorphic sex chromosomes in two flavors—old, undifferentiated sex chromosomes or newly-evolved sex chromosomes, due to a recent transition or failure to degenerate (Adolfsson and Ellegran, 2013; Furman et al. 2020; Kostmann et al. 2021). Since transitions can be frequent and difficult to identify, most transitions likely remain unknown. Empiricists now rely on recent advances in DNA sequencing and advanced cytogenetic methods to identify and characterize the evolution of homomorphic sex chromosomes (Augstenová et al. 2018; Gamble et al. 2015; 2017; Keating et al. 2020; 2021; Nielsen et al. 2018, 2019a, 2019b, 2020; Pan et al. 2019; 2021a; Rovatsos et al. 2019; Sidhom et al. 2020).

To date, high-throughput DNA sequencing technologies have largely confirmed early predictions that groups possessing homomorphic sex chromosomes frequently undergo transitions in sex chromosome systems (Augstenová et al. 2018; Blaser et al. 2013; Bull, 1983; Gamble et al. 2015a; Hundt et al. 2019; Jeffries et al. 2018; Kottler et al. 2020; Ogata et al. 2007; Uno et al. 2008). With only scant high-quality genomic resources available for many vertebrate groups, transitions have only been broadly estimated using changes in patterns in heterogamety, i.e. XY to ZW or vice versa (Gamble et al. 2015a). There are a few exceptions to this where, by chance, sex-linked genes can be identified and successfully mapped to a distant reference genome in order to identify a linkage group (e.g. Keating et al. 2020; Nielsen et al. 2019a; 2020). However, estimating the total number of SDS transitions from changes in heterogamety likely underestimates the true number of turnovers by a large margin (Gamble et al. 2015a; Jeffries et al. 2018). Indeed, current hypotheses suggest that homomorphic sex chromosomes may be maintained by frequent SDS turnovers that are more difficult to detect, *cis*-transitions, e.g. XY to new XY (Augstenová et al. 2018; Bachtrog et al. 2014; Blaser et al. 2014; Jeffries et al. 2018). To successfully recognize these more difficult to identify transitions and test hypotheses regarding sex chromosome turnover, we need high-quality reference genomes within groups that possess homomorphic sex chromosomes (Stöck et al. 2021).

Geckos (infraorder Gekkota) are a speciose clade of lizards that represent more than 1/3 of the known transitions in SDS’s within squamate reptiles (Gamble et al. 2015a; Gamble et al. 2018). Ancestrally, geckos possessed temperature-dependent sex determination (TSD) and have since undergone more than 25 transitions between TSD, XY, and ZW systems that are (generally) homomorphic (Gamble et al. 2015a; Pokorna and Kratochvíl L, 2009; Rovatsos et al. 2019). Although an extremely useful model system to study sex chromosomes, geckos currently have no chromosome-level reference genomes available to estimate linkage information for sex chromosome turnovers in geckos (Hara et al. 2018; Liu et al. 2015; Xiong et al. 2016). Thus, previous work in most gecko groups has been restricted to characterizing patterns of occurrence (which species are XY or ZW) without the ability to test hypotheses about sex chromosome conservation and turnover (although there are some exceptions, e.g. Keating et al. 2020; Nielsen et al. 2019a; Rovatsos et al. 2019; 2021). Indeed, until now this has also been the case for the charismatic, Neotropical geckos of the genus *Sphaerodactylus* (Figure 1).

**Figure 1:**
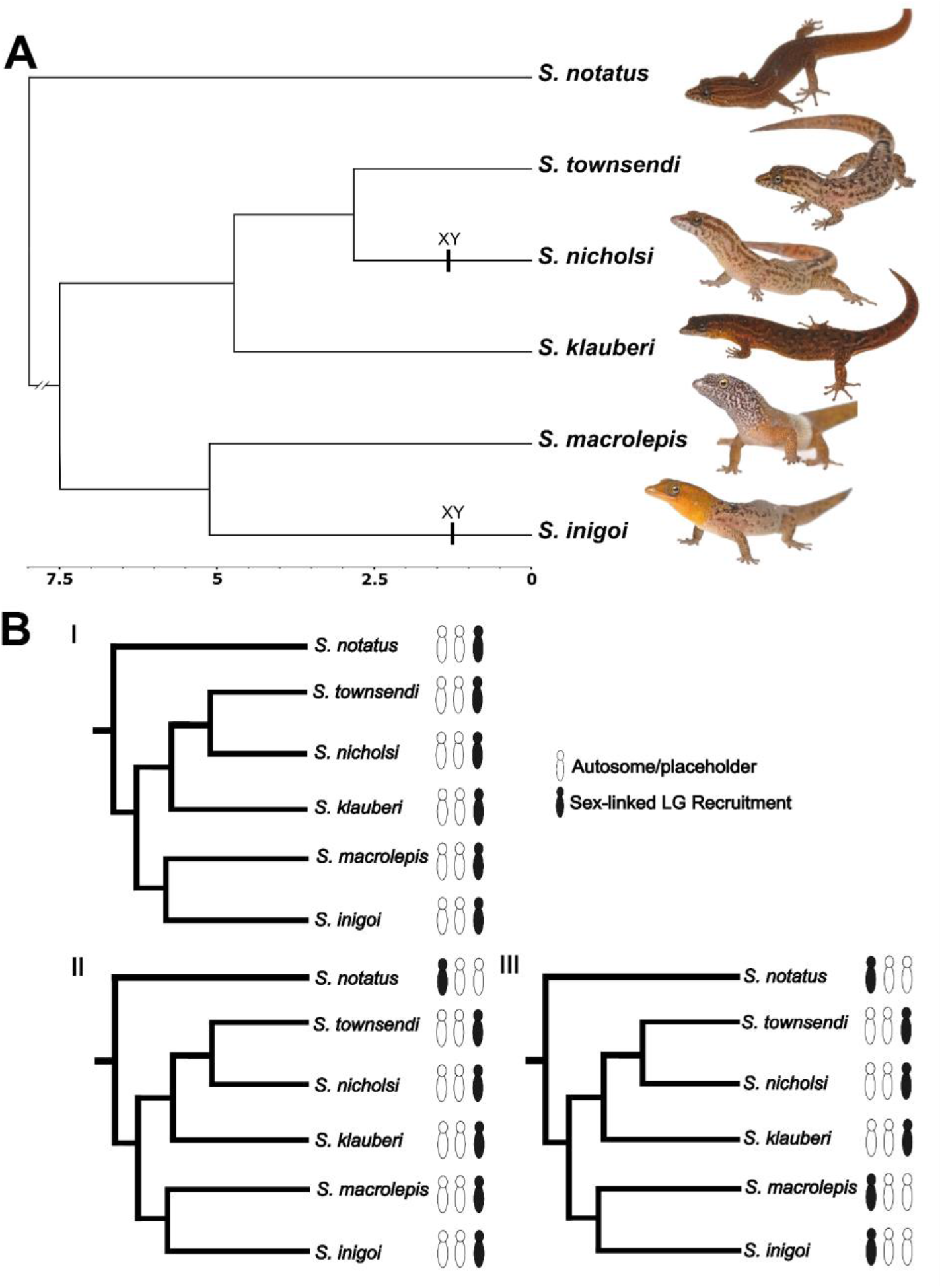
Overview of the study system and predictions of sex chromosome evolution in *Sphaerodactylus* geckos. (A) Time-calibrated phylogenetic tree (Daza et al. 2019) providing an evolutionary framework for *Sphaerodactylus* geckos from within the Puerto Rican Bank and an outgroup (*S. notatus*) with previously identified sex chromosome sex systems appended to tips (Gamble et al. 2015). (B) Three predictions for the observations of sex chromosome evolution in this group: (i) an ancestral XY system that has been conserved across all *Sphaerodactylus*; (ii) a conserved XY system within Puerto Rican *Sphaerodactylus*, but not across other sampled *Sphaerodactylus* species; and (iii) sex chromosome turnover among Puerto Rican *Sphaerodactylus* species.

The gecko family Sphaerodactylidae comprises 13 enigmatic genera distributed across 5 continents and a diversity of environments, yet only 4 genera have any information regarding sex chromosomes (reviewed in Gamble et al. 2018). To review these 4 genera briefly, karyotypes of male and female *Euleptes europaea* suggest an XY system with an unknown linkage group (Gornung et al. 2013). Recently, a conserved ZW system was discovered across the Caribbean genus *Aristelliger*, syntenic with *Gallus* chromosome 2 (Keating et al. 2020). Gamble et al. (2018) found an XY system in the northern South American *Gonatodes ferrugineus*, albeit with an unknown linkage group. Lastly, XY systems were discovered in *Sphaerodactylus nicholsi* and *Sphaerodactylus inigoi* (both native to the Puerto Rican Bank), also with unknown linkage groups (Gamble et al. 2015a). Taken together, these results suggest a high diversity of sex chromosome systems within Sphaerodactylidae and likely many more will be uncovered. However, the glaring deficiency—not knowing the linkage groups in most taxa—hampers our development of a broader understanding of sex chromosome evolution in this group. Therefore, the logical next step in diagnosing the diversity of sex chromosomes across sphaerodactylids is to begin assigning linkage groups to species with known SDS’s and their close relatives.

To begin addressing how sex chromosomes evolved in *Sphaerodactylus* geckos, we sequenced and assembled the first chromosome-scale gecko genome, Townsend’s least gecko (*Sphaerodactylus townsendi*) from Puerto Rico, and examined patterns of sex chromosome conservation and turnover in a small number of related *Sphaerodactylus* species. We chose to focus on the Puerto Rican *Sphaerodactylus* because we know more about their SDS systems than most other genera in the family (Gamble et al. 2015a; 2018) and a recent phylogenetic analysis provides a robust evolutionary framework (Daza et al. 2019; Pinto et al. 2019a). We generated a set of predictions for potential scenarios of sex chromosome evolution based on our previously identified knowledge of sex chromosomes within the group (outlined in Figure 1). We correlated these hypotheses with generalizable predictions regarding sex chromosome transitions in *Sphaerodactylus*: (i) an ancestral XY system has been conserved across sampled *Sphaerodactylus* species; (ii) a conserved XY system has been conserved across Puerto Rican *Sphaerodactylus*, but not maintained in other *Sphaerodactylus* clades, represented here by *S. notatus*; and (iii) sex chromosome systems have high rates of turnover across *Sphaerodactylus* taxa with distinct sex chromosome systems among the sampled Puerto Rican species.

To test these predictions, we collected a patchwork of genomic data from six *Sphaerodactylus* species (five from the Puerto Rican radiation and one outgroup, *S. notatus*, native to south Florida and the northern Caribbean) to accompany the new *S. townsendi* reference genome. This sampling represents <5% of described *Sphaerodactylus* species (6 of 107 species – Uetz et al. 2021). The data included in this study were: restriction-site associated DNA sequencing (RADseq); RNA sequencing (RNAseq); and whole-genome sequencing (re-sequencing). We used these data to definitively identify and confirm the sex chromosome linkage group in a subset of these species (*S. townsendi*, *S. nicholsi*, *S. inigoi*, and *S. notatus*). Then, we used a preliminary dataset generated from additional taxa (*S. klauberi* and *S. macrolepis*) to extrapolate from these more well-substantiated species to detect conserved patterns on the sex chromosomes. We identified multiple *cis*-transitions within *Sphaerodactylus* XY systems and report that these transitions are utilizing two linkage groups whose syntenic regions in chicken (*Gallus gallus*) were previously unknown to act as sex chromosomes in other tetrapods. Moreover, we begin to gauge the dynamic nature of sex chromosome evolution in *Sphaerodactylus*, which in turn may provide insight into the sex chromosome evolution of other underrepresented taxa with frequent sex chromosome transitions across the tree of life.

## Methods

### Data generation

We generated a high-quality reference genome for a male *S. townsendi* (indiv. TG3544 [male]) collected in Playa de Ponce, Puerto Rico (17.96439, -66.61387). Genome assembly combined linked-read sequencing (10X genomics), chromatin-contact sequencing (Hi-C), nanopore long-read sequencing, and whole-genome re- sequencing (WGS) using paired Illumina reads. For linked-read and nanopore long read sequencing, we extracted high molecular-weight (HMW) DNA from blood and liver tissue of one *S. townsendi* (TG3544 [male]) using a published DNA extraction protocol designed for low input (Pinto et al. 2021). For re-sequencing and all other DNA-related experiments described herein, we used Qiagen DNeasy^®^ DNA extractions of tail or liver tissues.

For the reference genome sequencing, we generated and sequenced a single 10X Chromium^®^ library (*S. townsendi* TG3544 [male]) across 2 lanes of Illumina^®^ HiSeqX (HudsonAlpha^®^ Institute for Biotechnology, Huntsville, AL); proximally-ligated input DNA from blood and liver tissue in-house (*S. townsendi* TG3718 [male]) using the Arima-HiC kit (Arima Genomics^®^, San Diego, CA USA) and sequenced it as a 400- 600bp insert Illumina^®^ library using the NEBNext Ultra II^®^ Library Preparation kit (New England Biolabs^®^ [NEB], Ipswich, MA USA) on an Illumina^®^ NovaSeq lane (Novogene^®^, Davis, CA USA); we generated 2 nanopore sequencing libraries (Oxford Nanopore^®^ Technologies [ONT], Oxford, UK) using the Ligation Sequencing Kit (SQK-LSK109) of HMW DNA (indiv. TG3544) and sequenced each library on its own flowcell (FLO- MINSP6) to completion (∼60 hours) on a single MinION^®^ device (MIN-101B); lastly, we made and sequenced a 400-600bp insert re-sequencing library on an Illumina^®^ HiSeqX (Psomagen^®^, Rockville, MD USA).

For reference genome annotation, we conducted additional RNA sequencing in *S. townsendi*. RNA sequencing (RNAseq) methods are described thoroughly by Pinto et al. (2019b); briefly, we extracted RNA from flash-frozen tissues stored at -80°C in Trizol^®^ reagent and generated sequencing libraries using the KAPA^®^ Stranded mRNA-Seq Kit for Illumina^®^ Platforms (KR0960 [v5.17]). We deep-sequenced RNAseq libraries from a whole head from a male (TG3467) and a whole embryo, 11 days post-oviposition (dpo) of unknown sex (TG3715), which were sequenced on an Illumina^®^ HiSeqX (Psomagen^®^, Rockville, MD USA). For downstream sex chromosome analyses (see below), we sequenced additional RNAseq libraries from whole heads (males and females) of *S. macrolepis* and *S. inigoi* preserved in RNAlater. These libraries were sequenced using paired-end reads (125-bp) on an Illumina^®^ HiSeq2500 (Medical College of Wisconsin, Milwaukee, WI USA).

To identify and explore the sex-linked regions of the genome, we generated additional whole-genome re-sequencing data for 1M/1F of *S. townsendi*, *S. nicholsi*, *S. klauberi*, and *S. notatus*. Additionally, we acquired population-level RADseq data for multiple males and females of *S. townsendi*, *S. nicholsi*, *S. inigoi*, and *S. notatus*. For whole genome re-sequencing data, we generated Illumina libraries for each individual using the NEBNext Ultra II kit (New England Biolabs). For RADseq data, we followed a modified protocol from Etter et al. (2011) as outlined in Gamble et al. (2015a). Libraries were pooled and sequenced using paired-end 100-bp or 150-bp reads on an Illumina^®^ HiSeq2000 at the University of Minnesota Genomics Center (Minneapolis, MN) or an Illumina^®^ HiSeqX at Psomagen^®^. In sum, our final dataset for assessing sex chromosome dynamics in *Sphaerodactylus* included: re-sequencing [1M, 1F] of *S. townsendi*, *S. nicholsi*, *S. klauberi*, *S. notatus,* and a single *S. macrolepis* male; RADseq data for *S. townsendi* [7M, 7F], *S. nicholsi* [6M, 6F], *S. inigoi* [7M, 9F] (from Gamble et al. 2015a), *S. notatus* [8M, 7F]; and RNAseq data from *S. macrolepis* [2M, 2F] and *S. inigoi* [2M, 2F]. The four RADseq species contained representative samples from across their known range. These data sources are summarized in Table 1.

**Table 1:**
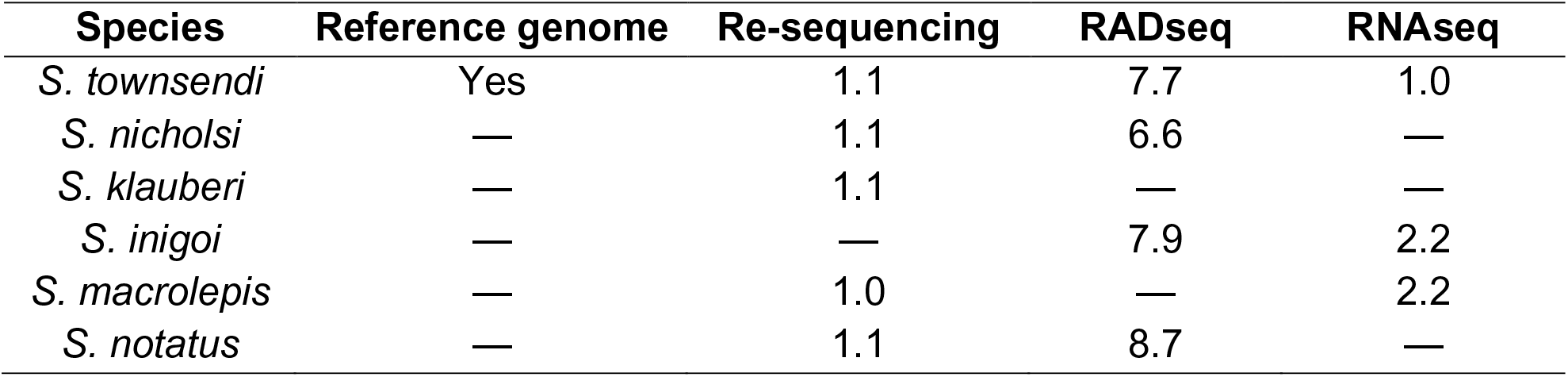
Table tracking the available data for each species used in this study. Notation in each cell refers to males and females (M.F).

### Transcriptome Assembly

We quality and adapter trimmed our RNAseq reads using Trim Galore!, filtered PCR duplicates using bbmap, and subsampled 50,000,000 PE reads for each tissue using seqtk. In an isolated docker computing environment (Merkel, 2014), we normalized cleaned reads and assembled *de novo* transcriptomes for each tissue using Trinity [v2.8.4] (Grabherr et al. 2011) in the *De novo* RNAseq Assembly Pipeline (DRAP) [v1.92] (Cabau et al. 2017). For *S. townsendi*, we generated both a ‘head’ and ‘embryo’ *de novo* assembly and combined them using the runMeta function in DRAP. An in-depth description of the utility of DRAP in the production of high-quality transcriptome assemblies can be found elsewhere (Cabau et al. 2017; Pinto et al. 2019b).

### Reference assembly, annotation, and characterization

We used a 6-part, iterative assembly approach to integrate the five different sequencing experiments (outlined in Table 2). In an effort to make these genome assembly efforts reproducible across platforms, all genome assembly steps—except for the initial SuperNova assembly (conducted at HudsonAlpha^®^) and three steps conducted in docker environments (details below)—were conducted in conda virtual environments that contained the following versions of these programs (in alphabetical order): ARCS [v1.1.1] (Yeo et al. 2018), assembly-stats [v1.0.1], bamtools [v2.5.1] (Barnett et al. 2011), BBmap [v38.79] (Bushnell, 2014), bcftools [v1.9] (Li, 2011), bedtools [v2.29.2] (Quinlan and Hall, 2010), diamond [v0.9.14] (Buchfink et al. 2015), freebayes [v1.3.2] (Garrison and Marth, 2012), HiSat2 [v2.1] (Kim et al. 2019), merqury [v1.3.0] (Rhie et al. 2020); minimap2 [v2.17] (Li, 2018), mosdepth [v0.2.6] (Pedersen and Quinlan, 2018), parallel [v20200322] (Tange, 2018), picard tools [v2.22], pixy [v1.1.1] (Korunes and Samuk, 2021), sambamba [v0.7.1] (Tarasov et al. 2015), samtools [v1.6] (Li et al. 2009), seqkit [v0.12] (Shen et al. 2016), seqtk [v1.3] (https://github.com/lh3/seqtk), STACKS [v2.3] (Catchen et al. 2013), Tigmint [v1.1.2] (Jackman et al. 2018), TGS-GapCloser [v1.0.1] (Xu et al. 2020), Trim Galore! [v0.5], and vcftools [v0.1.15] (Danecek et al. 2011).

**Table 2:** Tracking contiguity of the genome assembly across versions using 4 common metrics: Scaffold N50, size of the smallest scaffold comprising the largest 50% of the assembly; Scaffold L50 number of scaffolds comprising the largest 50% of the genome; Scaffolds, Total number of scaffolds comprising the full assembly; Size, The approximate number of base pairs in the assembly. BUSCO – Percent complete Core Vertebrate Genes (CVG).

| Assembly | Step | N50 | L50 | Scaffolds | Size | BUSCO |
| --- | --- | --- | --- | --- | --- | --- |
| v1.1 | SuperNova | 12,629,056 | 37 | 58,149 | 2.0 Gb | 93.6% |
| v1.2 | Tigmint | 6,460,730 | 69 | 59,469 | 2.0 Gb | 93.6% |
| v1.3 | ARCS | 7,457,274 | 57 | 58,603 | 2.0 Gb | 93.6% |
| v1.4 | TGS-GapCloser | 7,468,733 | 57 | 58,603 | 2.0 Gb | 94.9% |
| v1.5 | NextPolish | 7,605,248 | 57 | 58,603 | 2.0 Gb | 95.3% |
| v1.6 | 3D-DNA | 126,215,344 | 7 | 56,114 | 2.0 Gb | 95.7% |
| v1.7 | Redundancy-filter | 134,006,883 | 6 | 32,127 | 1.9 Gb | 95.7% |
| v1.8-v2.1 | +10kb cutoff | 134,006,883 | 6 | 1,823 | 1.8 Gb | 95.7% |

To assemble the reference genome from sequence data, we generated an initial assembly using SuperNova [v2.1.1] (Weisenfeld et al. 2017) using 80% our total 10X sequencing reads [assembly v1.1]. To improve this assembly, we broke misassemblies accumulated during the assembly process using Tigmint [assembly v1.2] and re-scaffolded with 100% of our 10X reads using ARCS [assembly v1.3]. Next, we incorporated the quality-filtered ONT reads (total reads = 435,394; total bp = 6,565,554,881; mean read length = 15,079.6; largest/smallest read = 162,107/1,001) to fill gaps in the genome using TGS-GapCloser [assembly v1.4]. Then, we combined Illumina data with ONT data to polish the genome using NextPolish [v1.3.1] (Hu et al. 2020) [assembly v1.5]. We broke and re-scaffolded the polished assembly using 2 iterations of 3D-DNA [v201008] (Dudchenko et al. 2017), which yielded 17 chromosome-scale scaffolds with no apparent large-scale misassemblies [assembly v1.6]. We visualized the final HiC contact map for misassemblies and with no large-scale misassemblies visible, we removed only small ‘blemishes’ from the contact map using Juicebox Assembly Tools [v1.11] (Durand et al. 2016). We removed duplicate assembled regions by mapping smaller assembled regions to the 17 chromosome-level scaffolds using RaGOO [v1.11] (Alonge et al. 2019) and removing scaffolds with high grouping confidence scores (i.e. 1.0) [assembly v1.7]. Lastly, to facilitate genome annotation, we removed scaffolds to a minimum length of 10Kb [assembly v1.8].

To functionally annotate the genome assembly, we used the Funannotate pipeline [v1.5.0] (Palmer, 2018) in an isolated docker computing environment (Merkel, 2014). Briefly, Funannotate provides a pipeline to soft-mask the assembly (https://github.com/Dfam-consortium/RepeatModeler) and predict gene models using both curated databases (Simão et al. 2015) and custom transcriptomic data (Haas et al. 2008; Hoff et al. 2015; Keller et al. 2011). To facilitate genome annotation, we provided transcriptomic data in the form of our aforementioned *de novo* meta transcriptome assembly. These files were then incorporated directly into the funannotate pipeline to inform the annotation process. The final annotated genome assembly was recoded to be submitted to GenBank as “MPM_Stown_v2.2”.

To assess the completeness and quality of the reference genome and *de novo* transcriptome, we employed metrics that query the assemblies for highly-conserved orthologous proteins. We used Benchmarking Universal Single-Copy Orthologs (BUSCO) [v5.1.2] (Simão et al. 2015), implemented on the gVolante web server [v2.0.0] (Nishimura et al. 2017), to query multiple databases of conserved orthologs: Core Vertebrate Genes (CVG) and tetrapoda_odb10. We calculated these metrics at each stage of genome assembly [assembly v1.1-v1.8] in its completeness as Supplemental Table 1 and present a subset of this information in Table 2. We also calculated completeness and quality metrics with kmers of our Illumina WGS using merqury.

### Sex chromosome identification and comparative genomics

#### WGS + RNAseq

We mapped WGS data to the genome using minimap2 and RNAseq data using hisat2. For WGS, we quantified per-individual read depth in 500Kb windows using mosdepth. We normalized each sample by its median read depth before calculating the male/female read depth in R [v3.6.2] (R Core Team, 2016). Importantly, for all species with WGS data, we identified no differences in read depth between males and females, which suggested that analyses examining sequence differences in this region would be successful. We called SNPs for WGS using freebayes to generate an ‘all-sites’ vcf file and calculated pi in 500Kb windowed using pixy. For RNAseq data, we called SNPs separately using freebayes to include only variable sites and calculated pi in 500Kb windowed using vcftools.

#### RADseq

Restriction site-associated DNA sequencing (RADseq) has been shown to be an essential tool for the identification of sex chromosome systems in species lacking heteromorphic sex chromosomes (e.g. Gamble et al. 2015a; 2018; Keating et al. 2020; Nielsen et al. 2019; 2020; Pan et al. 2021a). The expectation is that the only genomic region that should contain sex-specific RADtags are the non-recombining regions of the Y/W chromosomes (Gamble and Zarkower, 2014; Gamble, 2016). We can interpret areas with an abundance of mapped sex-specific RADtags as regions within the non-recombining region of the sex chromosomes. When analyzed alone, RADseq can identify sex chromosome systems but can say nothing of sex chromosome linkage or the size of the non-recombining region in the focal taxon (Fowler and Buonaccorsi, 2016; Gamble et al. 2015a; 2017; 2018; Hundt et al. 2019; Nielsen et al. 2019a; 2020). However, when analyzed in conjunction with a reference genome, we can both map these sex-specific RADtags to identify the linkage group and analyze the sequences in this region by calling SNPs from the raw data (Gamble, 2016; Pan et al. 2019). Thus, we can use both methods to confirm a region is sex-specific by looking for coincident locations of male/female differences across species.

### Reference-free analyses

We identified sex-specific RAD loci and their gametologous counterparts using the published RADtools pipeline plus a custom perl script (Gamble et al. 2015a; Nielsen et al. 2019b). For posterity, we validated a subset of *S. townsendi* RADtags as Y-linked via PCR (Figure 2); primer pairs S70_8.05_F1/R1 [[5’- CTTGTCACTTTTAGTGGGCACTG-3’/5’-GGATGCACGTTGTTGAACAAAAC-3’]] and S272_192_F2/R1 [5’-TTCAAAGCAAGAGATGTTCAGCG-3’/5’-GATCCTGGAATACGGMACCATGA-3’] (Figure 3), while those in *S. nicholsi* and *S. inigoi* were validated previously (Gamble et al. 2015).

**Figure 2:**
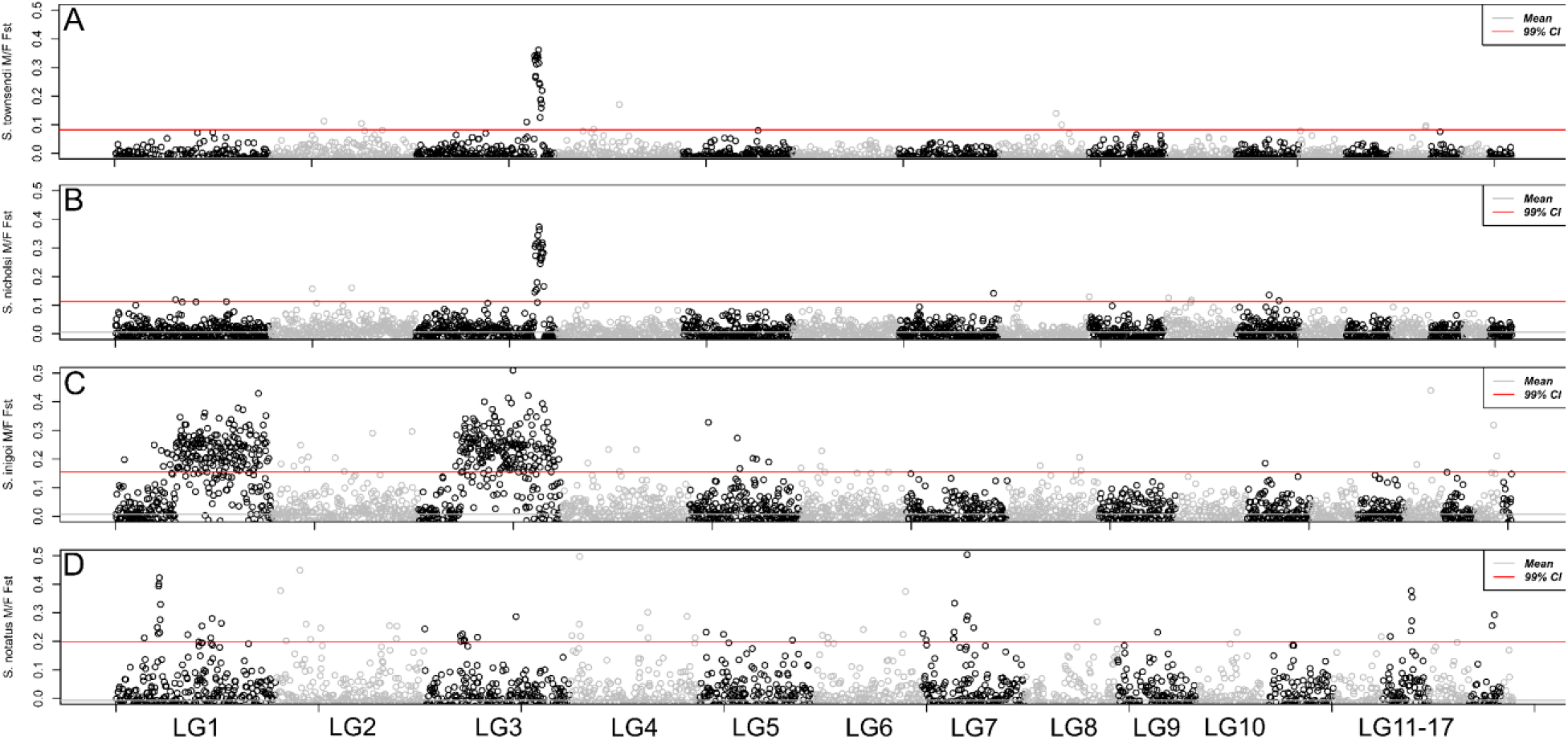
Whole-genome M/F F_ST_ scan in 500Kb windows using RADseq data for 4 taxa: (A) *S. townsendi*, (B) *S. nicholsi*, (C) *S. inigoi*, and (D) *S.notatus*.

**Figure 3:**
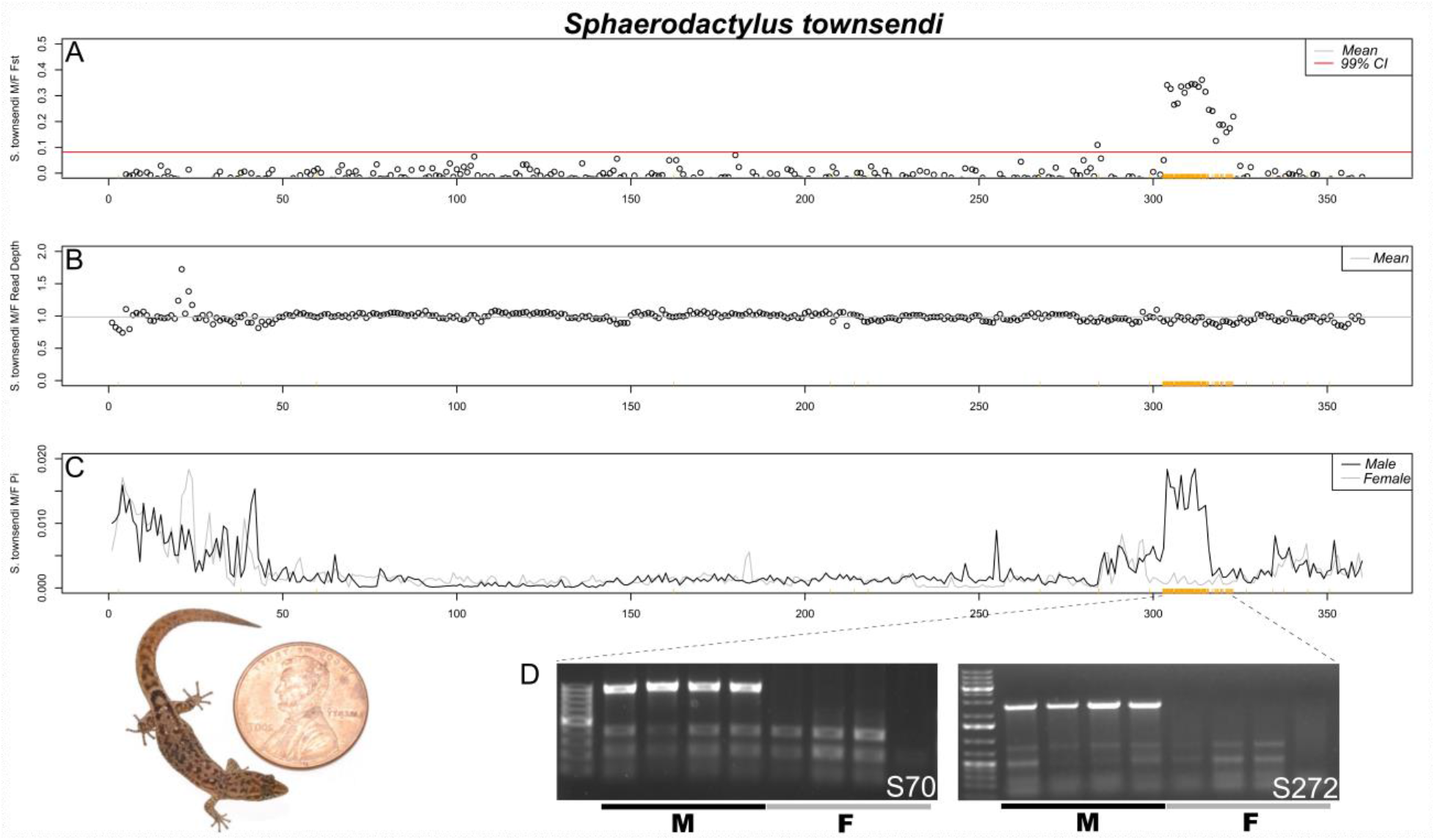
Confirmation of the *Sphaerodactylus townsendi* sex chromosome on LG3. (A) RADseq M/F F_ST_ scan in 500kb windows (zoomed in on LG3 from Figure 2); (B) M/F read depth differences across the length of LG3; (C) Male and female nucleotide diversity (π) along LG3. The same set of male-specific RADtags mapped to LG3 are denoted by orange ticks along the bottom of each graph (same in each panel). (D) Gel images from a subset of these markers illustrate that they are located on the Y chromosome. Picture of an adult male *S. townsendi* scaled with a penny, USA currency (diameter = 19.05 mm).

### Reference-assisted analyses

We mapped RADseq reads to the genome using minimap2 and used refmap.pl pipeline in STACKS to call SNPs separately for each species. We calculated male/female F_ST_ across the genome in 500Kb windows using vcftools and mapped the sex-specific RAD loci identified using the RADtools pipeline to the genome (Weir and Cockerham, 1984). We expected that each dataset would converge on specific areas in the genome (high M/F F_ST_ and many sex-specific markers added).

### Genome synteny and characterization

As this is the first chromosome-scale gecko reference genome assembled, we conducted a few analyses to characterize it relative to other reptile genomes.

Specifically, we identified the syntenic regions of the *S. townsendi* genome across 4 high-quality genomes available on Ensembl for reptiles: green anole (*Anolis carolinensis*, Acar2.0; Alföldi et al. 2011), Indian cobra (*Naja naja*, Nanav5; Suryamohan et al. 2020), common wall lizard (*Podarcis muralis*, PodMur_1.0; Andrade et al. 2019), and domestic chicken (*Gallus gallus*, GRCg6a). We identified syntenic regions in *S. townsendi* with these other reptile taxa using MCScanX (Wang et al. 2012). To visualize MCScanX synteny results, we generated synteny plots using SynVisio software (https://github.com/kiranbandi/synvisio). In addition, we explored GC content within *Podarcis*, *Anolis*, and *Naja* in 500Kb windows using python script (*slidingwindow_gc_content.py*) from Schield et al. (2019).

## Results

### Genome Characterization

We recovered 17 chromosome-length scaffolds for the *S. townsendi* reference genome. The best *a priori* estimate of haploid chromosome number in *Sphaerodactylus townsendi* is n=17—identified from the only three *Sphaerodactylus* karyotypes that currently exist, to our knowledge—the closely-related species: *S. ariasae*, *S. plummeri*, and *S. streptophorus* (presented here in Supplemental Figure 1). The total GC content was 46.0% (±11.1%) and we masked 44.47% of the genome modeled as repetitive DNA. Qualitatively, the assembly maintained high sequence accuracy and was largely complete. Our BUSCO score calculated against 5310 conserved tetrapod orthologs (tetrapoda_odb10) was 88.3% complete. Breaking this number down further, the assembly contained 87.6% single-copy orthologs, 0.7% duplicated ortholog copies, 3.9% fragmented copies, and 7.8% missing gene copies. When examining a subset of core vertebrate genes (CVG) with BUSCO, our assembly maintained a score of 95.7%. Similarly, we calculated 89.5% completeness value using our *S. townsendi* re- sequencing data with merqury.

We compared synteny maps with three other reptile species: chicken (*Gallus gallus*), green anole (*Anolis carolinensis*), and wall lizard (*Podarcis muralis*) and the information from the physically mapped *Gekko hokouensis* genome (Srikulnath et al. 2015). Naja was omitted from the table due to its collinearity with *Anolis* macrochromosomes. Most linkage groups (chromosome-scale scaffolds) maintained a one-to-one relationship with *P. muralis* chromosomes and maintained the known syntenic configurations in *G. hokouensis* (Table 3).

**Table 3:** A key to navigate synteny across largest fragments of the reference genome assembly relative to *Anolis, Podarcis*, and *Gallus*, according to the *Gekko hokouensis* (*Gekko*) physical mapping. Scaffolds were called if linkage groups described by Srikulnath et al. (2015) were corroborated by syntenic mapping to *Anolis*, *Podarcis*, and/or *Gallus*. Note that the snake (*Naja*) was omitted due to its collinearity with *Anolis* genome. *** indicates changes in annotated chicken chromosomes making up the linkage group from that reported by Srikulnath et al. (2015) from ‘21 and 25’ to ‘22 and 24’.

| *Sphaerodactylus townsendi* | *Anolis carolinensis* | *Podarcis muralis* | *Gallus gallus* | *Gekko hokouensis* |
| --- | --- | --- | --- | --- |
| LG1 | 1q | 3 | 3 | 1p |
| LG2 | 1p | 1 | 5,7 | 2 |
| LG3 (XY) | 2q | 2 | 12,13,16,18,30,33 | 1q |
| LG4 | 3q | 4 | 1q,14 | 13 |
| LG5 | 4q | 6,18 | 8,26,28 | 3 |
| LG6 | 5p | 10 | 1p,23 | 14 |
| LG7 | 2p | 11,17 | ZW | ZW |
| LG8 | 3p | 5,14 | 6,9 | 15 |
| LG9 | 4p | 7 | 2q | unplaced |
| LG10 | micro | 9 | 4q | 7 |
| LG11 | 6q | 12 | 2p,27 | 8 |
| LG12 | micro | 15,ZW | 17,22,24*** | 9 |
| LG13 | micro | 16,ZW | 4p,15 | 11 |
| LG14 | 4 | 8 | 11 | unplaced |
| LG15 | 6p | 13 | 27 | 12 |
| LG16 | micro | 8 | 21 | unplaced |
| LG17 | micro | 14 | 10 | unplaced |

### Sex chromosome identification and description

Across species with whole-genome re-sequencing data (WGS) for both a male and female, we observed no differences in read depth between the sexes (Figure 3; additional species not shown). Indeed, since read mapping did not differ between the sexes, we could successfully call SNPs and analyze sequence differences between the sexes. Thus, we called and analyzed SNPs for each of our datasets: WGS, RNAseq, and RADseq.

For species with RADseq data from multiple males and females, we identified a list of sex-specific RADtags using the Gamble et al. (2015) pipeline. For all species, we identified an excess of confirmed male-specific RADtags: *S. townsendi* (M = 431/F = 0), *S. nicholsi* (M = 186/F = 11), *S. inigoi* (M = 157/F = 0), and *S. notatus* (M = 21/F = 2). Previous work had validated a subset of these male-specific markers as Y-linked in *S. nicholsi* and *S. inigoi* using PCR (Gamble et al (2015). The majority of male-specific RADtags identified in each species, mapped to a small number of linkage groups in the *S. townsendi* genome: *S. townsendi* (LG3—87%), *S. nicholsi* (LG3—86%), *S. inigoi* (LG1—46%; LG3—51%), and *S. notatus* (LG1—62%). In concert, when examining male/female F_ST_, we observed a single, solitary peak of elevated F_ST_ in three species *S. townsendi* (LG3), *S. nicholsi* (LG3), *S. notatus* (LG1) (Figure 3), while *S. inigoi* presented two regions of elevated F_ST_ spanning both LG1 and LG3. The genomic regions where the sex-specific RADtags mapped overlapped with regions of elevated M/F F_ST_ values (LG3, Figure 4; LG1, Figure 5).

**Figure 4:**
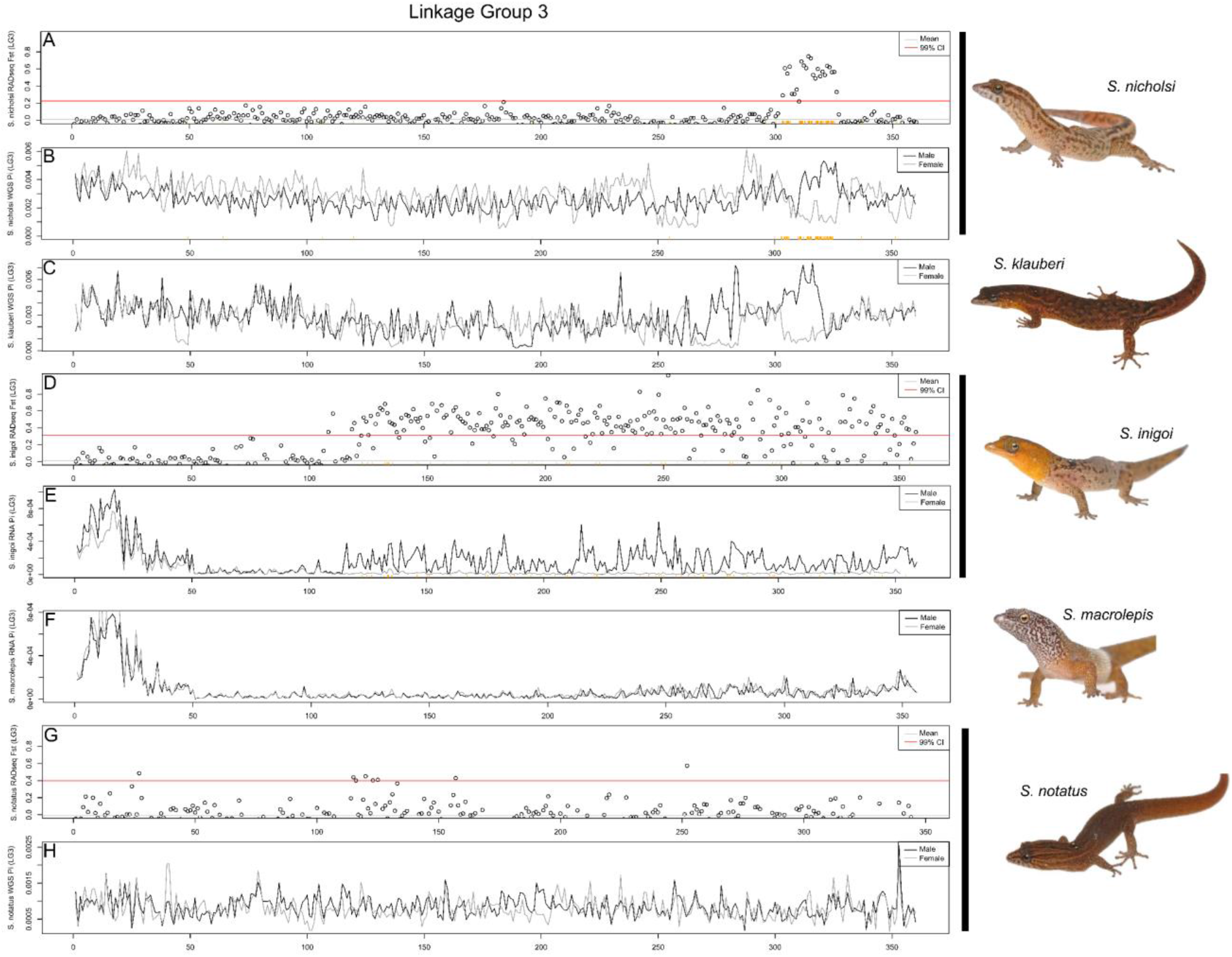
Comparative genomics of the *S. townsendi* sex chromosome (LG3) across multiple *Sphaerodactylus* species in 500kb windows. (A-B) *S. nicholsi* M/F F_ST_ values (RADseq) and M+F nucleotide diversity (WGS), respectively; (C) *S. klauberi* M+F nucleotide diversity (WGS); (D-E) *S. inigoi* M/F F_ST_ values (RADseq) and M+F nucleotide diversity (RNAseq), respectively; (F) *S. macrolepis* M+F nucleotide diversity (RNAseq); (G-H) *S. notatus* M/F F_ST_ values (RADseq) and M+F nucleotide diversity (WGS), respectively. Sex-specific RADtags mapped to *S. nicholsi* (A-B) and *S. inigoi* (D-E) along the X axis (orange ticks). Note: slight shifts on the X-axis are due to the differences in programs used to calculate values, i.e. WGS used pixy, while RADseq and RNAseq used vcftools.

**Figure 5:**
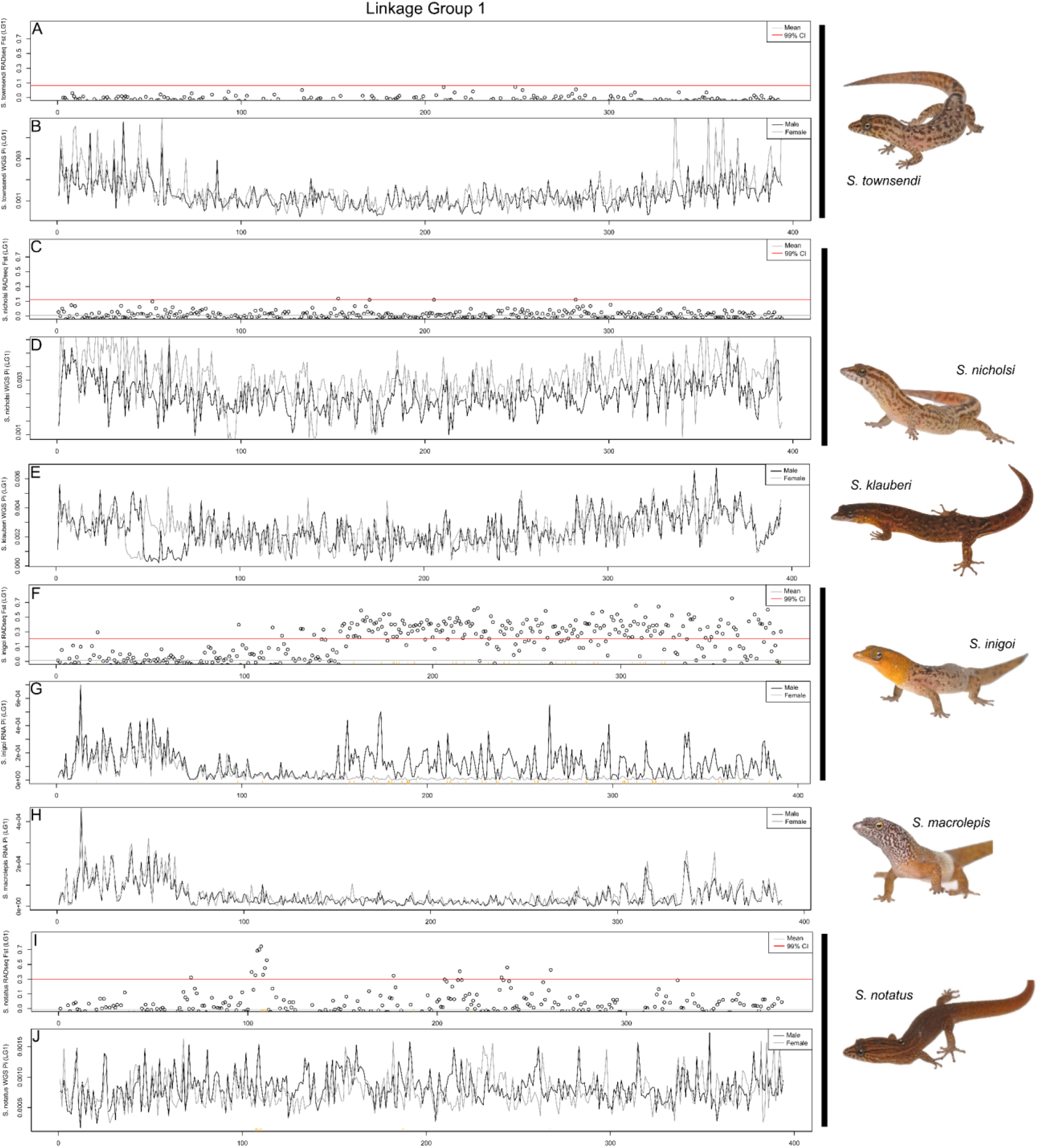
Comparative genomics of the *S. notatus* sex chromosome (LG1) across multiple *Sphaerodactylus* species in 500kb windows. (A-B) *S. townsendi* M/F F_ST_ values (RADseq) and M+F nucleotide diversity (WGS), respectively; (C-D) *S. nicholsi* M/F F_ST_ values (RADseq) and M+F nucleotide diversity (WGS), respectively; (E) *S. klauberi* M+F nucleotide diversity (WGS); (F-G) *S. inigoi* M/F F_ST_ values (RADseq) and M+F nucleotide diversity (RNAseq), respectively; (H) *S. macrolepis* M+F nucleotide diversity (RNAseq); (I-J) *S. notatus* M/F F_ST_ values (RADseq) and M+F nucleotide diversity (WGS), respectively. Sex-specific RADtags mapped to *S. inigoi* (F-G) and *S. notatus* (I-J) along the X axis (orange ticks). Note: slight shifts on the X- axis are due to the differences in programs used to calculate values, i.e. WGS used pixy, while RADseq and RNAseq used vcftools.

**Figure 6:**
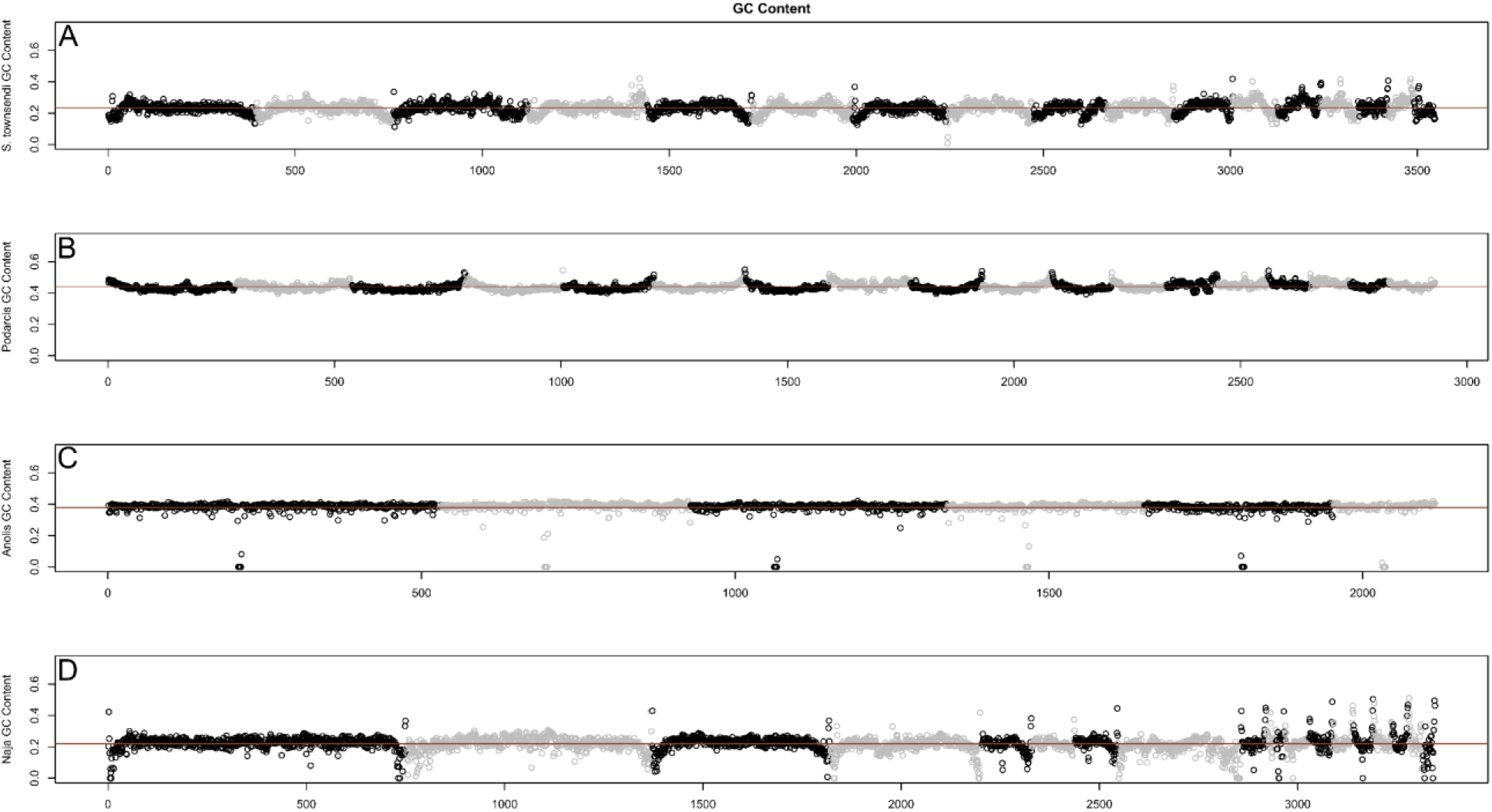
Genome-wide patterns of GC content across representative squamate taxa, orange line representing the genomic mean. Broadly, pattern of GC content appears most similar, both in chromosome patterns and mean per-window GC content (∼0.2), between (A) *Sphaerodactylus* and (D) *Naja*. Both (B) *Podarcis* and (C) *Anolis* have a considerably higher mean per-window GC (∼0.4), and *Podarcis* shows an inverse pattern to *Sphaerodactylus* and *Naja* in that GC goes up at the tips of chromosomes instead of down. We believe that the *Anolis* patterns here are less informative in this regard as the sequencing method employed is not directly comparable to the other 3 genomes.

After identifying the non-recombining region of the sex chromosomes in each species with RADseq, we used the WGS and RNAseq datasets to further characterize and corroborate these regions. We calculated nucleotide diversity (π) across the sex chromosomes. In recently evolved non-recombining regions where both X and Y reads map; we expect increased nucleotide diversity in males due to the increase in heterozygosity in this region relative to the rest of the genome (Schield et al. 2019). We confirmed that this was indeed the case in species where we had already identified the sex chromosomes using RADseq, i.e. *S. townsendi*, *S. nicholsi*, *S. inigoi*, and *S. notatus* (Figure 3a, 3c, 4b-e, 5f-g, 5i-j; Supplemental Figure 3 and 4). Then, we looked to species without available RADseq data, i.e. *S. klauberi* and *S. macrolepis*. We observed an increase in male π in *S. klauberi* WGS data at the same location in the sister species (*S. townsendi* and *S. nicholsi*), suggesting a conserved XY system in this clade (Figure 4c). However, we saw no such elevation in π, nor F_ST_, in *S. macrolepis* RNAseq data, indicating a lack of a non-recombining region in males on this linkage group (Figure 4f, 5h; Supplemental Figure 5).

Lastly, we used the RNAseq data in *S. townsendi* and *S. inigoi* to explore whether both X and Y alleles are both being expressed in this system, essentially using these data as another reduced-representation genomic dataset (similar to RADseq).

Indeed, for *S. townsendi* and *S. inigoi*, we scanned each for genomic signatures identified in RADseq and WGS data (e.g. Figure 2). In *S. townsendi*, we identified a peak in male nucleotide diversity that coincides with the identified SDR on LG3 (Supplemental Figure 2). In *S. inigoi*, we observed the same patterns in the RNAseq data as seen in RADseq data for both LG1 and LG3 calculating both F_ST_ (Supplemental Figures 3 and 4) and nucleotide diversity (Figures 4d-e and 5f-g). However, in *S. macrolepis*, for whom we also had RNAseq data, we saw no differences between males and females on either linkage group coinciding with the SDRs identified in this study when examining F_ST_ (Supplemental Figure 5) or nucleotide diversity (Figures 4f and 5h); nor did we see any elevation in nucleotide diversity in the single male WGS data (not shown). Thus, we are as of yet unable to identify the sex chromosome linkage group in *S. macrolepis*, but these data suggest that it likely not the same system as either *S. townsendi* or *S. inigoi*.

We examined synteny across the genome to construct a quick-reference synteny table correlating each *S. townsendi* linkage group with their syntenic regions in *Podarcis*, *Anolis*, and *Gallus* (Table 3). Of note, the Indian cobra (*Naja naja*) was omitted from the table as its macrochromosome were collinear with *Anolis*. We used these correlations to approximate the locations of these linkage groups in the physically mapped *Gekko hokouensis* genome (Srikulnath et al. 2015). More specifically, for the sex chromosomes we present a fine-scale synteny analysis comparing the sex chromosome linkage groups identified here with their counterparts in *Podarcis*, *Anolis*, and *Gallus* (Supplemental Figure 6). We identified that most *Sphaerodactylus* linkage groups are represented in other species as a single syntenic block (e.g. *Podarcis* and *Gallus* macrochromosomes), while others are whole chromosome arms (*Anolis* macrochromosomes) or made up of many smaller linkage groups in other more distantly-related lineages (i.e. *Gallus* microchromosomes). This information provides a simple reference for future work investigating genome synteny in geckos.

## Discussion

### Reference Genome Description

The final genome assembly of *Sphaerodactylus townsendi* achieved chromosome-level status (Table 3). At the time of submission, this is the first such assembly in a gecko and one of only a handful of high-quality assemblies in squamate reptiles. Other publicly-available chromosome-level squamate assemblies include those from the Indian cobra (*Naja naja;* Suryamohan et al. 2020) and prairie rattlesnake (*Crotalus viridis*; Schield et al. 2019), as well as the physically-mapped green anole (*Anolis carolinsensis*) genome (Alfoldi et al. 2011), common wall lizard (*Podarcis muralis*; Andrade et al. 2019), and Komodo dragon (*Varanus komodoensis*; Lind et al. 2019) genomes, with more being sequenced, assembled, and published on a regular basis. A non-exhaustive list of the currently available Lepidosaur reference genomes is provided in Supplemental Table 3.

### Sex chromosome evolution in Sphaerodactylus

Across our sampled taxa, we found that five out of six *Sphaerodactylus* species have XY sex chromosomes and the sixth, *S. macrolepis*, remains unknown. Among the taxa with an identified sex chromosome system, three maintain a conserved XY system encompassing (presumably) a single stratum of the sex-determining region (SDR) on LG3 (*S. townsendi*, *S. nicholsi*, and *S. klauberi*). Our outgroup, *S. notatus*, possesses a distinct sex chromosome system located on LG1, which rejects a hypothesis of a conserved XY system across the sampled *Sphaerodactylus* species (Figure 1b: Hypotheses I). *S. inigoi* maintains a sex chromosome system that includes both LG1 and LG3, likely due to chromosomal fusion. The *S. inigoi* XY system is extremely large and encompasses most of LG1 and LG3, including the SDR region of *S. townsendi* on LG3 but excluding the SDR of *S. notatus* on LG1. Thus, we cannot reject the hypothesis that *S. townsendi* and *S. inigoi* inherited their sex chromosome system from their most-recent common ancestor (MRCA) (e.g. Figure 1b: Hypotheses II or III). Notably, the sex chromosome system in *S. macrolepis* remains unknown, deviating slightly from our specific predictions. However, finding no evidence in *S. macrolepis* of the stark patterns of sex linkage in our *S. macrolepis* data (although sparse) leads us to predict that there may have been a transition within this lineage that requires additional work to elucidate, but we will not discuss this null result further here. We divide the rest of this section into two parts based on two potential interpretations of these findings: (1) LG3 was the sex chromosome in the MRCA of *S. townsendi* and *S. inigoi*, i.e. these systems are homologous (Figure 1b: Hypothesis II); or (2) the observations of LG3 as a sex chromosome in *S. townsendi* and *S. inigoi* represent independent recruitment events of LG3 as a sex chromosome (Figure 1b: Hypothesis III).

#### (1) Hypothesis II: A conserved XY system within Puerto Rican Sphaerodactylus, but not across other Sphaerodactylus species

After two species diverge, the non-recombining region of their sex chromosomes (that were inherited from a common ancestor) can fluctuate between species, both in genomic location (e.g. addition of evolutionary strata) and level of sequence degeneration (Graves, 2008; Lahn and Page, 1999). In the case of *Sphaerodactylus* from Puerto Rico—*S. townsendi*, *S. nicholsi*, *S. klauberi* (herein the “*S. townsendi* group”), and *S. inigoi*—all have a sex-linked LG3. However, the non-recombining region of the Y in *S. inigoi* entirely encompasses the SDR of the *S. townsendi* group. The most parsimonious way to generate this pattern is a single origin of the LG3 linkage group as a sex chromosome. Alternatively, if the SDR identified in the *S. townsendi* group was present in the MRCA of *S. townsendi* and *S. inigoi*—having remained largely static in *S. townsendi* but expanded greatly in *S. inigoi*—we might expect to see an overall increase of sex-specific markers or F_ST_ values located in that region in *S. inigoi* (indicating an older stratum), and/or conserved male-specific RAD markers on the Y chromosomes of each species. However, we see none of these lines of evidence, suggesting that this SDR may not have been present in the MRCA of *S. townsendi* and *S. inigoi*.

#### (2) Hypothesis III: Sex chromosome systems have high rates of turnover in Sphaerodactylus

Although it is certainly possible for a sex chromosome system on LG3 to have been inherited from a MRCA, it is equally likely that LG3 (similarly to LG1 in *S. inigoi* and *S. notatus*) was recruited independently into a sex-determining role. While there are numerous examples for recent *cis-* (XY to XY) and *trans*- (e.g. XY to ZW) transitions in sex chromosomes to different linkage groups at shallow scales (e.g. Jeffries et al. 2018; Tao et al. 2020), there are far fewer confirmed examples of *cis*-transitions to the same linkage group. Empirical examples of *trans*-transitions to the same linkage group in other systems have emerged in recent literature, such as: the Japanese wrinkled frog (*Glandirana rugosa*) possessing independently-derived XY and ZW systems on the same linkage group with two independent derivations of the ZW system accompanied by lineage-specific W-degradation (Ogata et al. 2003; 2007; Miura et al. 2011); *Xiphophorus maculatus* chromosome 21 (Xma21) has been recruited multiple times within the genus *Xiphophorus* (Franchini et al. 2018). However, *cis*-transitions to the same linkage group have only been identified in recent years within ranid frogs (Jeffries et al. 2018), stickleback fishes (independent derivations of an XY system on LG12; Ross et al. 2009), and possibly also multiple times within *Xiphophorus* fishes (M. Schartl *pers. comm.*). Thus, although there are fewer known *cis*-transitions to the same linkage group, they are also much harder to diagnose (as witnessed by this study), which we posit is the most parsimonious reason for this discrepancy in the current literature.

In the vertebrate literature, some linkage groups have been recruited as sex chromosomes multiple times, while others have remained unutilized (Graves and Peichel, 2010; Kratochvíl et al 2021; O’Meally et al. 2012). For example, the syntenic regions with the bird ZW system have been independently recruited as a sex chromosome in both a turtle (*Staurotypus triporcatus*; XY) and two geckos (*Gekko hokouensis*; ZW and *Phyllodactylus wirshingi*; ZW) (summarized in Nielsen et al. 2019a). In sphaerodactylids, the only linkage group previously identified as a sex chromosome linkage group was the ZW system in *Aristelliger* (*Anolis* 6/*Gallus* 2; Keating et al. 2020). Within *Sphaerodactylus*, this is the first identified use of the *S. townsendi* LG1 (syntenic with *Gallus* 3) or LG3 (specifically sections of the chromosome syntenic with *Gallus* 18/30/33; Table 3) in geckos (Augstenová et al. 2021). Beyond geckos, to our knowledge it is the first time in any tetrapod that the syntenic regions of *Gallus* 3 and 30/33 have been recruited as a sex chromosome (Kratochvíl et al. 2021). *Sphaerodactylus townsendi* LG3 has only been found as a partial component (i.e. *Gallus* chromosome 18, not including *Gallus* 30/33) of the sex chromosome linkage group in one other species, the ZW system of the night lizard *Xantusia henshawi* (Nielsen et al. 2020). Thus, no other vertebrate group (to our knowledge) has recruited either of these linkage groups as a sex chromosome. One possible reason that these syntenic regions have not been previously identified as a sex chromosome linkage group is an apparent lack of ‘usual suspect’ sex determining genes (Graves and Peichel, 2010; Herpin and Schartl, 2015).

In many vertebrate groups where empiricists have identified the master sex determining gene (MSD), a list of commonly-recruited MSDs have been identified (i.e. the ‘usual suspects’; Dor et al. 2019; Herpin and Schartl, 2015). The same genes have been co-opted to function as the MSD in other groups, including *Dmrt1* in birds, a frog (*Xenopus laevis*), and some medaka fish (members of the *Oryzias latipes* group); *Sox3* in placental mammals and other medaka (members of the *Oryzias celebensis* and *O. javanicus* groups); and *Amh* in tilapia and pike and other fishes (Li et al. 2015; Myosho et al. 2015; Pan et al. 2019; and see Pan et al. 2021b for recent review). Interestingly, we identified one candidate MSD located on LG3 from the list of ’usual suspects’ (SRY-Box Transcription Factor 9; *Sox9*), but, at least in the *S. townsendi* group, it is located outside of the hypothesized non-recombining region and no paralogous copies of *Sox9* are found elsewhere in the genome. Thus, we suggest that its value as a candidate sex-determining gene in this system is limited. However, the presence of this gene (and potentially other genes) may pre-dispose this linkage group as a repeatably-evolving sex chromosome that may yet be identified in other systems (Graves and Peichel, 2010; O’Meally et al. 2012), or perhaps these species utilize a previously unidentified MSD, lending support to the null hypothesis that any linkage group may act a sex chromosome given the proper selective pressures (e.g. Hodgkin, 2002). We also identified no candidate MSD genes in the much smaller *S. notatus* SDR region on LG1.

### Genome Architecture and Synteny

Qualitative patterns of windowed GC content were most-similar between the *Sphaerodactylus* and *Naja* despite not being more closely related to each other than other sampled taxa. Interestingly, both geckos and snakes are ancestrally nocturnal (Gamble et al. 2015b; Pinto et al. 2019c; Simões et al. 2016), and it is plausible the genome-wide decrease in per-window GC content resulted in independent losses of highly thermo-stable DNA in both lineages (Fullerton et al. 2001). Alternatively, this could also be lineage-specific to *Sphaerodactylus* (Scantlebury et al. 2011). The global patterns of GC content being more similar between *Anolis* and *Podarcis* than that of *Sphaerodactylus* is surprising in that *Sphaerodactylus* and *Podarcis* are two of the very few squamate reptiles that do not possess microchromosomes (Olmo et al. 1990; Srikulnath, 2013). Indeed, contrasting patterns of genome-wide patterns of GC content (and potentially other indicators of genome organization) differ to such an extent could be explained by two independent origins of macrochromosome-only karyotypes.

Alternatively, these patterns could be explained by changes in recombination landscape between taxa (Charlesworth, 1994) or related to the presence/absence of isochores (Eyre-Walker et al 2001). These ideas should be further tested with multiple chromosome-level genome assemblies across geckos, snakes, and additional squamates.

### Future Directions

In recent years, much has been learned about vertebrate sex chromosome evolution. Just within geckos, we have expanded our knowledge of sex chromosome systems (SCS) exponentially (see Augstenová et al. 2021 and Gamble et al. 2015a). In sphaerodactylids, we have discovered three distinct sex chromosome linkage groups within two genera (Keating et al. 2020; *this study*) and at least two other XY systems that currently lack linkage information (Gamble et al. 2018; Gornung et al. 2013). The identification of sex chromosome linkage groups is fast becoming more feasible as new reference genomes become available, as well as new tools that permit the more critical analysis of mechanisms of sex determination and sexual differentiation, both practically and financially (Rasys et al. 2019; Stöck et al. 2021). Thus, the sprightly sphaerodactyls are poised to become a potent model system for genomic research. We here point out two potentially worthwhile research avenues.

First, future work focusing on unsampled species both nested within our focal taxa (e.g. *S. macrolepis* and others), closely-related outgroup taxa (e.g. *S. roosevelti*), as well as more-distant relatives, could help develop a clear hypothesis for when and how these newly identified sex chromosome linkage groups were recruited within this genus. For example, a closer look at the sister species of *S. inigoi*, *S. grandisquamis*, may provide insight into the timing of the putative chromosomal fusion we hypothesize in *S. inigoi* and identifying biological correlates of sex chromosome transitions (e.g. population bottlenecks during island colonization; Daza et al. 2019) may help better estimate the total number of SCS transitions in other groups. Research including *S. roosevelti* and other more distantly related species will illuminate whether LG3 is a sex chromosome linkage group that was inherited from a common ancestor or independently derived between the two Puerto Rican clades (the clades containing *S. inigoi* + *S. grandisquamis* and *S. townsendi* + *S. klauberi*; see Daza et al. 2019). The above questions specifically focusing on ‘when did these SCS evolve?’ logically lead to more intricate questions regarding SCS stability and their influence within the sex determination signaling cascade.

Second, sphaerodactylids display an impressive diversity of morphological characteristics, such as body size and sexual dichromatism. Indeed, it has been posited that sexual dichromatism has evolved repeatedly within *Sphaerodactylus* (Daza et al. 2019; Regalado, 2015). Coincidentally, one such loss of dichromatism is hypothesized between the sister clades containing the dichromatic *S. inigoi* + *S. macrolepis* and the monochromatic *S. townsendi* + *S. klauberi*. If this dichromaticism is influenced by sex chromosomes in *S. inigoi* (encompassing almost 2 entire chromosomes)—and that degenerated system is ancestral—the loss of the *S. inigoi* system in the *S. townsendi* clade could have been selected for to relieve predation pressures, to resolve sexual conflict, etc… (Stöck et al. 2011; van Doorn and Kirkpatrick, 2007).

## Conclusions

We presented data and analyses of the sex chromosomes for a small percentage of the known taxonomic diversity within *Sphaerodactylus* geckos. Within this small subset of species, our analyses reject the hypothesis that there is a conserved SCS maintained across *Sphaerodactylus* geckos. We identified and further characterized between 2 and 4 *cis*-transitions between species with XY sex chromosome systems.

These two newly identified sex chromosome linkage groups are syntenic with regions that have not previously been characterized as sex chromosomes in an amniote: LG1 (syntenic with *Gallus* 3) and LG3 (syntenic regions with *Gallus* 18/30/33). We posit that the recruitment of *S. townsendi* LG3 as a sex chromosome in *S. townsendi* and *S. inigoi* is likely independent. We reviewed the data for and against multiple recruitments of this chromosome between these taxa and suggest that a putative sex chromosome fusion in *S. inigoi* may correspond with the sex chromosome transition specific to this lineage. Overall, we show that present estimates of sex chromosome transitions (which are conservative, e.g. Augstenová et al. 2021) within gecko lizards are likely even more conservative than previously conceived, making geckos an even more essential model to study sex chromosome evolution (Gamble et al. 2015a; 2018; Nielsen et al. 2019a; Rovatsos et al. 2019). These results further build on a growing body of work indicating that gekkotan sex chromosome evolution is far more intricate than previously hypothesized and ready for further study.

## Data Availability

Sequencing reads generated for this study: 10X Chromium^®^ sequencing, long-read sequencing, Illumina^®^ re-sequencing, RNAseq, and RADseq are available on NCBI’s Sequence Read Archive (SRA) under project number PRJNA746057 and a full list of individual metadata is available in Supplemental Table 3. Assembled and annotated genome and transcriptomes are available on Figshare (https://doi.org/10.6084/m9.figshare.12291236) and the annotated genome will be available on NCBI after processing under genome accession XXXXXX000000000. All genomic computation took place on a custom-built, 24-core Intel Xeon^®^ and 128Gb RAM system running Ubuntu 16.04.

## Acknowledgements

Authors would like to thank J. Bernstein, M. “Toño” García, E. Glynne, N. Holovacs, J. Titus-McQuillan, C. Rivera, V. Rodriguez, D. Zarkower, E. Blumenthal, and D. DeFilippis for their respective contributions to the completion of this work. Animals were collected with permission of the Puerto Rican Departamento de Recursos Naturales y Ambientales (DRNA; under permits 2014-IC-042, 2013-IC-006, and 2016- IC-091). All experiments were carried out in accordance with Institutional Animal Care and Use Committee (IACUC) protocols at Marquette University (AR279 and AR288) and the University of Minnesota: 0810A50001 and 1108A03545. Specimens from Florida were collected by the permission of the Florida Fish and Wildlife Conservation Commission given to the Florida Museum of Natural History. Sociedad Ornitológica de la Hispaniola for assisting with logistics and the Ministerio de Medio Ambiente y Recursos Naturales for providing them with permits necessary for the collection and exportation of specimens in the Dominican Republic (0512-0515).

## Author Contributions

BJP designed study, directed fieldwork, conducted genome sequencing and later RADseq experiments, acquired funding, assembled and analyzed data, and wrote the manuscript; SEK constructed RNAseq libraries; SVN conducted fieldwork and constructed RADseq and RNAseq libraries; DPS conducted fieldwork; JDD conducted fieldwork and provided organismal expertise; TG assisted in design of study, conducted fieldwork and initial RADseq experiments, performed karyotyping, acquired funding, and oversaw the project. All authors read and approved the final manuscript.

